# Long-delay Learning in Intraoral Conditioned Taste Aversion

**DOI:** 10.1101/026500

**Authors:** Thomas A. Houpt, Jennifer A. Cassell, Stefanie McCormack, Bumsup Kwon, Gary Tiffany, Jeffrey Lorch

## Abstract

The cardinal feature of conditioned taste aversion (CTA) learning is the ability of animals to associate the taste or flavor of a food (the conditioned stimulus; CS) with a subsequent toxic effect (unconditioned stimulus; US), even if the toxicity occurs hours later, i.e. after a long delay. Two experiments were conducted which took advantage of the stimulus control afforded by intraoral catheterization to establish the parameters of long-delay learning in intraoral CTA. First, to determine the range of CS-US intervals which supports intraoral conditioning, rats received infusions of 5% sucrose paired with LiCl (76 mg/kg, ip) across a range of delays (0-6 h). Second, to determine the interaction of US dose and delay, rats were conditioned with sucrose paired with different doses of LiCl (19, 38 or 76 mg/kg) at several CS-US intervals (0, 10, or 60 min). Conditioning, assessed during a second infusion of sucrose at 48 h post-conditioning, was optimal at 10 min (although not significantly different at intervals between 0 and 60 min). Effectiveness declined at longer delays, such that CTA was not supported at intervals of 3h or greater. The dose-interval function suggested that an increased US can compensate for a longer CS-US interval. Low doses of LiCl induced a long-term CTA at 0-min (19 and 38 mg/kg) or 10-min delays (38 mg/kg), but were not sufficient to induce CTA at longer delays, which required the highest dose (76 mg/kg).

## 1. Introduction

The temporal characteristics of conditioned taste aversion (CTA) learning are similar in many aspects to those of other forms of classical conditioning. CTA requires the contingent pairing of a conditioned stimulus (the CS, a taste) with an unconditioned stimulus (the US, a toxin such as low-dose LiCl). CTA learning is not readily acquired after backwards conditioning (i.e. toxin presented before taste stimulus (Barker & Smith, 1974; Nachman, 1970).) CTA learning has short-term and long-term phases: the short-term phase (less than 6 h) is rapidly expressed within minutes after contingent taste-toxin pairing (Eckel & Ossenkopp, 1996; Houpt & Berlin, 1999; Spector, Breslin, & Grill, 1988), is labile after weak conditioning and decays within hours (Houpt & Berlin, 1999), and is protein-synthesis independent (Houpt & Berlin, 1999; Lamprecht, Hazvi, & Dudai, 1997). Likewise, the consolidation of long-term CTA learning (after 4-6 h) requires strong conditioning and protein synthesis (Houpt & Berlin, 1999; Rosenblum, Meiri, & Dudai, 1993; Tucker & Gibbs, 1976).

The cardinal feature of CTA learning, however, is the ability of animals to associate taste with a subsequent toxic effect, even if the toxicity occurs hours later, i.e. after a long delay (Garcia, McGowan, & Green, 1966; J. C. Smith & Roll, 1967). This is a unique property for simple associative learning: other forms of classical conditioning (e.g. fear conditioning or eye-blink conditioning) require overlap of the conditioned and unconditioned stimulus (Kimble & Reynolds, 1967); at most they tolerate delays of milliseconds to seconds between the offset of the CS and the onset of the US (the interstimulus interval, or ISI). Compared to these forms of conditioning, even the common CTA protocol, of pairing a taste stimulus (the CS) with a toxin (the US) after a delay of 15 min to 1 h after the taste, is remarkable. Yet, CTA acquisition has been reported even with CS-US intervals of 6-12 h (Burritt & Provenza, 1991; Kronberg, Muntifering, & Ayers, 1993; Revusky, 1968; J. C. Smith & Roll, 1967). Support of a long-delay is likely a beneficial trait: the ability to associate the flavor of a food with a delayed postingestive effect would be an obvious advantage to survival in an environment with potentially toxic or contaminated foods. It has even been suggested that CTA learning not only tolerates a delay, but that it is optimized for a long delay to match the time course of postingestive bacterial incubation or absorption of toxins (Schafe, Sollars, & Bernstein, 1995).

One method for inducing CTA employs intraoral catheters to delivery the CS directly into the rat’s mouth, thus obviating the need for voluntary food ingestion by the animal (Grill & Norgren, 1978). Intraoral conditioning has several advantages: it allows administration of a fixed gustatory stimulus with precise timing of onset and duration regardless of the animal’s motivation to ingest (Houpt, Philopena, Joh, & Smith, 1996); it facilitates evaluation of stimulus-induced orofacial motor behaviors (Grill & Berridge, 1985) and neural responses (Houpt, Philopena, Wessel, Joh, & Smith, 1994); and it separates consummatory responses from appetitive behaviors through bypass of the detection and approach phase of feeding (Wolgin & Wade, 1990). It has also been proposed that intraoral CTA learning, because it removes the operant responses underlying normal ingestion, reflects a more classical Pavlovian type of conditioning, with a greater dependence on the amygdala (Schafe, Thiele, & Bernstein, 1998).

In this study we took advantage of the stimulus control afforded by intraoral delivery to establish the parameters of long-delay learning in intraoral CTA. Two experiments were conducted. First, to determine the range of delays which supports intraoral conditioning, rats received intraoral infusions of 5% sucrose paired with LiCl injections (76 mg/kg) across a range of CS-US intervals (0-6 h). Second, to determine the interaction of US dose and delay, rats were conditioned with sucrose paired with different doses of LiCl (19,38 or 76 mg/kg) at several CS-US intervals (0, 10, or 60 min).

## 2. Methods

### 2.1 Animals and Intraoral Catheterization

Adult male Sprague-Dawley rats (Charles River) were individually housed at 25° C under a 12:12 light:dark cycle with ad lib access to Purina rodent chow and water except as noted. Under halothane anesthesia, rats were implanted with intraoral catheters made of PE-90 tubing that entered the mouth through the lateral check and were externalized on the dorsal surface between the scapulae, as described previously (Houpt & Berlin, 1999). Intraoral catheters were flushed daily with water to maintain patency. For intraoral infusions, rats were weighed and placed in a glass aquarium subdivided into 4 individual compartments by Plexiglas sheets. Conditioning was carried out 4-5 days after catheterization. Syringe pumps infused 5% sucrose into the mouth at a rate of 1 ml/min for 6 min. After the infusion, the intraoral catheter was flushed with distilled water, and the rat along with any feces were weighed again as a measure of consumption. Rats were returned to their home cages after infusion, where LiCl injections were administered.

### 2.2 Range of CS-US Intervals

Rats (n=71) were implanted with intraoral catheters as above. After overnight food deprivation, the rats were infused with 5% sucrose (the CS) over 6 min at 1 ml/min. Rats were injected with LiCl (0.15M, 76 mg/kg, i.p.; the US) at one of 9 time points relative to the start of the intraoral infusion (CS-US interval): 0, 10, 20, 30, 60, 120, 180, 240, and 360 min. (The 0-min injection was administered immediately before the start of the infusion). As a non-contingent control, additional rats were injected with LiCl at 180 min before the intraoral infusion (-180- min interval). Food and water were returned after the LiCl injections.

Rats were conditioned in 2 cohorts at 5 CS-US intervals each (0-60 min, and 60-360 min), and both cohorts included 60-min and -180-min groups. In total, 11 rats were conditioned at the -180-min interval, 12 rats were conditioned at the 60-min interval, and 6 rats were conditioned at each of the remaining 8 intervals.

The next day, rats were again overnight food-deprived. Approximately 48 h after the conditioning infusion of sucrose, long-term CTA expression was assessed. Rats were weighed and intraorally infused with 5% sucrose over 6 min at 1 ml/min, then weighed again immediately after the infusion to determine intraoral intake.

### 2.3 US Dose and CS-US Interval

Rats (n=72) were implanted with intraoral catheters as above. After overnight food deprivation, the rats were infused with 5% sucrose (the CS) over 6 min at 1 ml/min. Rats were injected with LiCl (the US) at one of 4 doses: 0, 19, 38, or 76 mg/kg made 0.3 Osm with NaCl (at 12 ml/kg i.p.), at one of 3 time points relative to the start of the intraoral infusion (CS-US interval): 0, 10, and 60 min. Six rats were conditioned at each dose and CS-US interval. Food and water were returned after the LiCl injections.

The next day, rats were again overnight food-deprived. Approximately 48 h after the conditioning infusion of sucrose, long-term CTA expression was assessed. Rats were weighed and intraorally infused with 5% sucrose over 6 min at 1 ml/min, then weighed again immediately after the infusion to determine intraoral intake.

### 2.4 Statistical Analysis

Data are presented as the mean ± standard error of the mean. T-tests, one- and two-way ANOVAs, and Tukey-Kramer HSD post hoc comparisons were performed using Xynk software (http://xynkapp.com). Exponential curve fitting was performed with Kaleidagraph software (Synergy Software).

## 3. Results

### 3.1 CS-US Intervals

On the day of conditioning, most rats consumed most of the intraoral infusion of 5% sucrose. However, there was a significant effect of treatment on CS intake on conditioning day (F(9,64) = 2.09, p < 0.05). Only the 0-min group (which received a LiCl injection immediately before the intraoral infusion) drank less sucrose compared to the -180-min group (3.7 ± 0.4 g vs. 5.7 ± 0.3 g, p< 0.05), although 0-min group intakes were not significantly lower than any of the other groups (mean 5.0 ± 0.2 g).

Intraoral intake of the CS at 48 h after conditioning was significantly affected by treatment (one-way ANOVA, F(9,64) = 8.95, p < 5e-08; see Figure 1). The non-contingent control (-180-min) group consumed almost all of the infused sucrose. Groups conditioned with a CS-US interval of 0 to 60 min had significantly reduced intake compared to the non-contingent control group. CS intakes in the 180-, 240-, and 360-min delay groups were significantly higher than the minimal intake of the 10-min delay group. Thus, intraoral conditioning with 5% sucrose and 76 mg/kg LiCl supports acquisition of a significant long-term CTA at CS-US intervals from 0 to 60 min, with an intermediate effect at 120 min.

**Figure 1.**
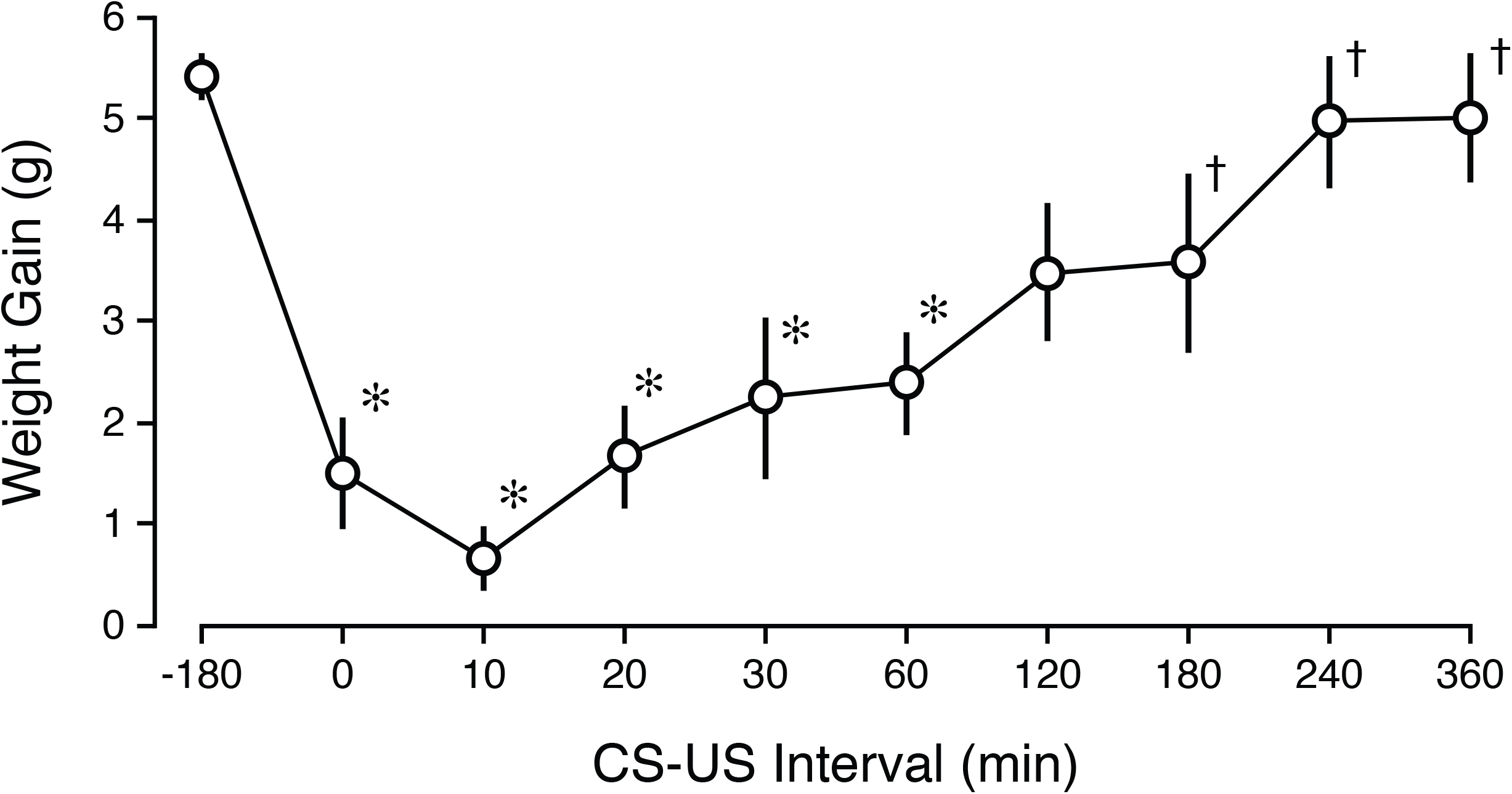
Long-delay CTA expression assessed with intraoral intake. Intraoral intake of 5% sucrose (6 ml / 6 min) after one pairing of intraoral sucrose and LiCl (0.15 M, 12 ml/kg) at CS-US delay intervals. n= 6-12 at each time point. * p < 0.05 vs. -180 min CS-US interval; † p < 0.05 vs 10 min CS-US interval.

The attenuation of CTA with increased CS-US interval suggests a decay of a “CS signal” across the ISI. We therefore fitted an exponential decay function to the 48-h intake (inverted to suppression from 6 g) of the 0-360 min groups (R[61] = 0.66, p < 0.001):

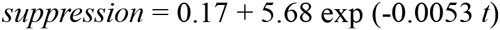

Based on this exponential decay function, the maximum suppression after conditioning would lower intake from 5.8 g (unconditioned) to 0.2 g (optimal conditioning). The calculated half-life of the “CS signal” is 131 min, with a mean lifetime of 189 min. Although the fitted decay function was statistically significant even with this small data set, the fitting would be, of course, more accurate with greater number and precision of data points.

### 3.2 US Dose and CS-US Interval

There was a significant interaction of LiCl Dose and CS-US interval on CS intake on conditioning day (two-way ANOVA, F(6,60) = 2.81, p < 0.05), such that rats which received 76 mg/kg LiCl immediately before the intraoral infusion of sucrose (0-min) had significantly lower intake compared to other groups (3.5 ± 0.5 g vs. 5.2 ± 0.1 g)

During CTA expression at 48 h after conditioning, there was an effect of LiCl Dose (F(3,60) = 39.00, p < 5e-14) and CS-US interval (F(2,60) = 13.39, p < 5e-05) on intraoral intake of the CS; the interaction of dose and interval approached significance (F(6,60) = 2.11, p = 0.065; see Figure 2). Compared to the high intake of the unconditioned control groups at 0 mg/ kg, CS intake was significantly lower i) at all doses of LiCl after a 0-min delay, ii) at 38 and 76 mg/kg LiCl after the 10-min delay, and iii) only at 76 mg/kg after the 60-min delay (p’s < 0.05). The groups that received 76 mg/kg LiCl after 0-min and 10-min delays also drank significantly less CS than groups that received 19 mg/kg after 10- and 60-min delays and 38 mg/kg after a 60- min delay.

**Figure 2.**
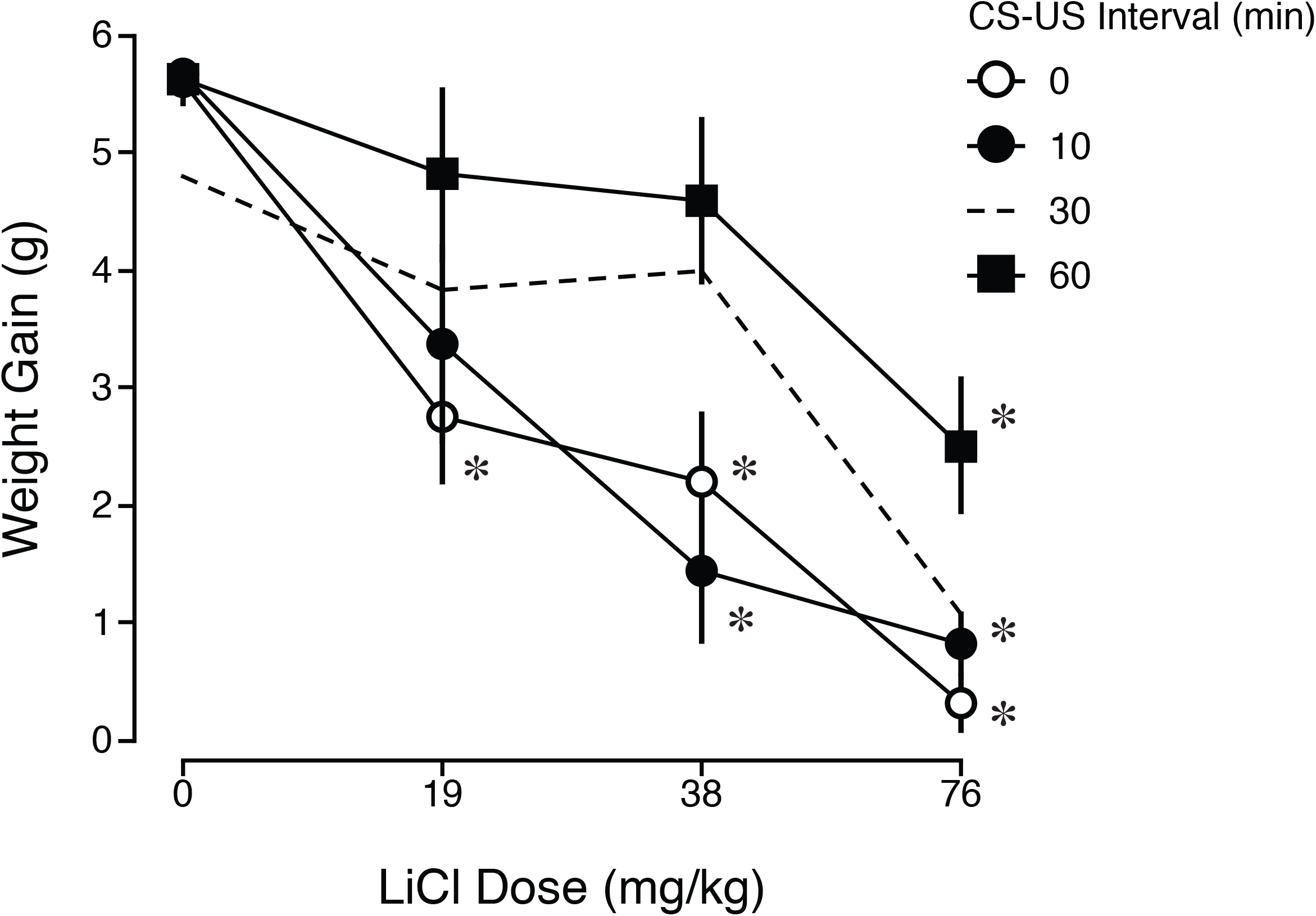
Interaction of CS-US interval and US dose in long-delay CTA expression. Intraoral intake of 5% sucrose (6 ml / 6 min) after one pairing of intraoral sucrose and LiCl at one of 4 doses (0, 19, 38 or 76 mg/kg) at one of 3 CS-US delay intervals: 0 (white circles), 10 (black circles), or 60 min (black diamonds). For comparison, data at 30-min CS-US interval (dashed line) is replotted from Houpt and Berlin 1999 (Houpt & Berlin, 1999) but was not included in the present statistical analysis. * p < 0.05 vs. 0 mg/kg dose at 0 min CS-US interval.

We note that these results are complementary to earlier published data collected in the same protocol with 19, 38, and 76 mg/kg LiCl at a 30-min CS-US interval (Houpt & Berlin, 1999), as replotted in Figure 2. The dose-response functions suggest that a stronger US permits consolidation of a CTA after longer CS-US intervals.

## 4. Discussion

These results add to earlier studies of long-delay learning in CTA. Long-delay learning was first discovered by Garcia et al. in 1966 (up to 3 h) (Garcia et al., 1966) and Smith et al. in 1967 (up to 12 h) (J. C. Smith & Roll, 1967). Many groups have demonstrated long-delay CTA learning, and there have been several systematic explorations of the limits of the CS-US interval in rats (Garcia et al., 1966; Kalat & Rozin, 1971; Nachman, 1970; Nachman & Jones, 1974; Revusky, 1968; Schafe et al., 1995; J. C. Smith & Roll, 1967; Wright, Foshee, & McCleary, 1971) and other species (Burritt & Provenza, 1991; Kronberg et al., 1993).

CTA induced by intraoral infusions also supports long-delay learning (Grill & Berridge, 1985; Grill & Norgren, 1978), and we here provide a parameterization of effective CS-US intervals using intraoral sucrose paired with LiCl. Conditioning was optimal at 10 min (although not significantly different at intervals between 0 and 60 min). Effectiveness declined at longer delays, such that CTA was not supported at intervals greater than 2 h. In comparison to other published CS-US interval curves, conditioning with intraoral CTA was not as robust as conditioning induced and measured with ad libitum CS intake (e.g. by 2-bottle tests which support delays of 6-12 h between comparable sucrose-LiCl pairings (J. C. Smith & Roll, 1967)). In effect, intraoral CTA requires a higher dose of LiCl to achieve the same magnitude as bottle conditioning (Limebeer & Parker, 2006; Wolgin & Wade, 1990). Intraoral and bottle CTA may engage different mechanisms of learning, such as classical vs. operant conditioning (Schafe et al., 1998); alternatively, the reflexive consummatory response to intraoral sucrose may simply be more resistant to learned changes compared to voluntary appetitive intake from a bottle (Limebeer & Parker, 2006).

Sufficient data points were examined to allow fitting of an exponential decay function to the CS-US interval curve. Exponential decay of a CS trace have often been proposed in theories of classical conditioning, for example modeling the dissipation of intracellular signals triggered by the CS (Balkenius & Morén, 1998; Gingrich & Byrne, 1987; Gluck & Thompson, 1987). Without a neurological model, it is impossible to interpret what process is decaying during CTA learning, but fitting a quantitative relationship may assist future correlations. Famously, Kalat and Rozin proposed a model in which the decay of CTA learning across long-delays was not due to the decay of a CS engram, but rather due to the simultaneous increase in “learned safety” to the CS flavor which inhibited CTA (Kalat & Rozin, 1973). However, there has been little recent work to explore their model, particularly at the neural or molecular level.

### US Contribution to Long-Delay Learning

By assessing CTA learning across multiple time points with multiple doses of the US, we demonstrated that a larger dose of LiCl can support CTA learning across a longer CS-US interval. The effect of US dose on 48-h expression may reveal an interaction with the process of CTA consolidation. Low doses of LiCl induced a long-term CTA (as measured at 48 h post-conditioning) at 0-min (19 and 38 mg/kg) or 10-min delays (38 mg/kg), but not at longer delays. However, these same doses of LiCl can support expression of a 30-min long-delay CTA in the short-term (as measured at < 6h h post-conditioning), during the protein synthesis-independent phase of CTA memory (Houpt & Berlin, 1999). Thus the apparent inability of low-doses of LiCl to support longer CS-US intervals, when measured at 48 h, may be masked by the lack of long-term memory consolidation. In fact, the non-genomic substrates of short-term CTA memory are sufficient to support at least 30-min CS-US intervals.

Earlier studies (Andrews & Braveman, 1975; Wright et al., 1971) have also consistently demonstrated that a stronger US permits learning at longer delays. These dose-response functions suggest that an increased US can compensate for the decay of a CS signal across the CS-US interval. This property might be reflected in neural sites of taste-toxin convergence.

## Conclusion

Despite the prominent role of long-delay learning in early CTA studies, its mechanism is still obscure. The intraoral infusion technique aids the exploration of the long-delay by providing precise temporal control of CS delivery. To further quantify the relative contributions of the gustatory CS and toxic US, a parallel study should be carried out with intraoral infusions of the CS at different concentrations or durations (Steinert, Infurna, Jardula, & Spear, 1979). In fact, a full isobolographic analysis would be required (i.e., treating CS and US as 2 drugs which act synergistically to induce CTA (Braude & Ginzburg, 1986)). Although the onset of CS stimulation and US effects can be controlled relatively precisely, the project is complicated by the need to match the intensity and duration of CS during conditioning and expression tests, and to account for the magnitude and duration of unconditioned responses to the US.

## References

Andrews, E. A., & Braveman, N. S. (1975). The combined effects of dosage level and interstimulus interval on the formation of one-trial poison-based aversions in rats. Animal Learn Behav, 3, 287–289.

Balkenius, C., & Morén, J. (1998). Computational models of classical conditioning: a comparative study. In R. Pfeifer, B. Blumberg, J.-A. Meyer, & S. W. Wilson (Eds.), From animals to animats 5:proceedings of the fifth international conference on simulation of adaptive behavior (pp. 348–353). Cambridge: MIT Press/Bradford Books.

Barker, L. M., & Smith, J. C. (1974). A comparison of taste aversions induced by radiation and lithium chloride in CS-US and US-CS paradigms. J. Comp. Physiol. Psychol., 87(4), 644–654.

Braude, M. C., & Ginzburg, H. M. (1986). Strategies for research on the interactions of drugs of abuse. NIDA Research Monograph, 68, 1–239.

Burritt, E. A., & Provenza, F. D. (1991). Ability of lambs to learn with a delay between food ingestion and consequences given meals containing novel and familiar foods. Appl Anim Behav Sci, 32(2-3), 179–189.

Eckel, L. A., & Ossenkopp, K. P. (1996). Area postrema mediates the formation of rapid, conditioned palatability shifts in lithium-treated rats. Behavioral Neuroscience, 110(1), 202– 212.

Garcia, J., McGowan, B., & Green, K. (1966). Learning with a prolonged delay of reinforcement. Psychon Sci, 5, 121–122.

Gingrich, K. J., & Byrne, J. H. (1987). Single-cell neuronal model for associative learning. Journal of Neurophysiology, 57(6), 1705–1715.

Gluck, M. A., & Thompson, R. F. (1987). Modeling the neural substrates of associative learning and memory: a computational approach. Psychological Review, 94(2), 176–191.

Grill, H. J., & Berridge, K. C. (1985). Taste reactivity as a measure of the neural control of palatability. Prog. Psychobio. Physiol. Psych., 11, 1–61.

Grill, H. J., & Norgren, R. (1978). Chronically decerebrate rats demonstrate satiation but not bait shyness. Science (New York, NY), 201(4352), 267–269.

Houpt, T. A., & Berlin, R. (1999). Rapid, labile, and protein synthesis-independent short-term memory in conditioned taste aversion. Learning & Memory (Cold Spring Harbor, NY), 6(1), 37–46.

Houpt, T. A., Philopena, J. M., Joh, T. H., & Smith, G. P. (1996). c-Fos induction in the rat nucleus of the solitary tract by intraoral quinine infusion depends on prior contingent pairing of quinine and lithium chloride. Physiology & Behavior, 60(6), 1535–1541.

Houpt, T. A., Philopena, J. M., Wessel, T. C., Joh, T. H., & Smith, G. P. (1994). Increased c-fos expression in nucleus of the solitary tract correlated with conditioned taste aversion to sucrose in rats. Neuroscience Letters, 172(1-2), 1–5.

Kalat, J. W., & Rozin, P. (1971). Role of interference in taste-aversion learning. J. Comp. Physiol. Psychol., 77(1), 53–58.

Kalat, J. W., & Rozin, P. (1973). “Learned safety” as a mechanism in long-delay taste-aversion learning in rats. J. Comp. Physiol. Psychol., 83(2), 198–207.

Kimble, G. A., & Reynolds, B. (1967). Eyelid conditioning as a function of the interval between conditioned and unconditioned stimuli. Foundations of Conditioning and Learning, Ed. G.a. Kimble, 279–287.

Kronberg, S. L., Muntifering, R. B., & Ayers, E. L. (1993). Feed aversion learning in cattle with delayed negative consequences. Journal of Animal Science, 71(7), 1767–1770.

Lamprecht, R., Hazvi, S., & Dudai, Y. (1997). cAMP response element-binding protein in the amygdala is required for long-but not short-term conditioned taste aversion memory. The Journal of Neuroscience : the Official Journal of the Society for Neuroscience, 17(21), 8443–8450.

Limebeer, C. L., & Parker, L. A. (2006). Effect of conditioning method and testing method on strength of lithium-induced taste aversion learning. Behavioral Neuroscience, 120(4), 963–969. http://doi.org/10.1037/0735-7044.120.4.963

Nachman, M. (1970). Learned taste and temperature aversions due to lithium chloride sickness after temporal delays. J. Comp. Physiol. Psychol., 73(1), 22–30.

Nachman, M., & Jones, D. R. (1974). Learned taste aversions over long delays in rats: the role of learned safety. J. Comp. Physiol. Psychol., 86, 949–956.

Revusky, S. H. (1968). Aversion to sucrose produced by contingent x-irradiation: temporal and dosage parameters. J. Comp. Physiol. Psychol., 65(1), 17–22.

Rosenblum, K., Meiri, N., & Dudai, Y. (1993). Taste memory: the role of protein synthesis in gustatory cortex. Behavioral and Neural Biology, 59(1), 49–56.

Schafe, G. E., Sollars, S. I., & Bernstein, I. L. (1995). The CS-US interval and taste aversion learning: a brief look. Behavioral Neuroscience, 109(4), 799–802.

Schafe, G. E., Thiele, T. E., & Bernstein, I. L. (1998). Conditioning method dramatically alters the role of amygdala in taste aversion learning. Learning & Memory (Cold Spring Harbor, NY), 5(6), 481–492.

Smith, J. C., & Roll, D. L. (1967). Trace conditioning with X-rays as an aversive stimulus. Psychon Sci, 9, 11–12.

Spector, A. C., Breslin, P., & Grill, H. J. (1988). Taste reactivity as a dependent measure of the rapid formation of conditioned taste aversion: a tool for the neural analysis of taste-visceral associations. Behavioral Neuroscience, 102(6), 942–952.

Steinert, P. A., Infurna, R. N., Jardula, M. F., & Spear, N. E. (1979). Effects of CS concentration on long-delay taste aversion learning in preweanling and adult rats. Behavioral and Neural Biology, 27(4), 487–502.

Tucker, A., & Gibbs, M. (1976). Cycloheximide-induced amnesia for taste aversion memory in rats. Pharmacology Biochemistry and Behavior, 4(2), 181–184.

Wolgin, D. L., & Wade, J. V. (1990). Effect of lithium chloride-induced aversion on appetitive and consummatory behavior. Behavioral Neuroscience, 104(3), 438–440.

Wright, W. E., Foshee, D. P., & McCleary, G. E. (1971). Comparison of taste aversion with various delays and cyclophosphamide dose levels. Psychon Sci, 22, 55–56.

